# Spatial distribution and research trend of illegal activities and the factors associated with wild mammal population declines in protected areas

**DOI:** 10.1101/2019.12.16.877944

**Authors:** Alfan A. Rija, Rob Critchlow, Chris D. Thomas, Colin M. Beale

**Author notes:** Correspondence author: Dr Alfan A. Rija.

## Abstract

Illegal activities are a persistent problem in many protected areas, but an overview of the extent of this problem and its impact is lacking. We review 35 years (1980-1914) of research across the globe to examine the spatial distribution of research and socio-ecological factors influencing population decline within protected areas under illegal activities pressure. From 92 papers reporting 1048 species/site combinations, more than 350 species comprising mammals, reptiles, birds, fishes and molluscs were reported to have been extracted illegally from 146 protected areas across four continents. Research in illegal activities has increased substantially during the review period but also shows strong taxonomic and geographic biases towards large wild mammals and African continent respectively, suggesting persistent poaching pressures on wild mammals in African protected areas. Population declines were most frequent i) where there was commercial poaching as opposed to subsistence poaching alone, ii) in countries with a low human development index particularly in strict protected areas and iii) for species with a body mass over 100 kg. Habitat loss associated with greater land use change had an additional significant impact on population decline, particularly in the less-strict categories (IUCN III-VI) of protected area across the continents. Overall, these findings provide evidence that illegal activities are most likely to cause species declines of large-bodied animals in protected areas in resource-poor countries regardless of protected area conservation status (i.e. IUCN category). Given the mounting pressures of illegal activities, additional conservation effort such as improving anti-poaching strategies and conservation resources in terms of improving funding and personnel directed at this problem is a growing priority.

## Introduction

Improving the spatial coverage of protected area (PA) network is increasingly viewed as a global biodiversity conservation priority (1, 2). The global land area with legal protection for conservation has increased from 3.5% in 1985 (3) to 13% recently (4) and looks set to increase further as countries aim to fulfil the Aichi target of protecting 17% of terrestrial land by 2020 (5). Despite this increase and the investment made in protected areas (6), biodiversity loss is perceived to be continuing even in PAs. For example, Craigie, Baillie (7) reported continent-wide population declines of large mammals across several of Africa’s protected areas. To allocate conservation resources efficiently, it is important to understand the scale and trends of illegal activities. Most studies, e.g. Mitchell (8), Wright, Zeballos (9), Leader-Williams, Albon (10) have focused on a single threat, on a single protected area type or region, and/or over short time periods. These provide crucial PA-specific data, but information on illegal activities and their impact at broader spatial and temporal scales is sparse, making tackling illegal activities difficult. Here we review the site-specific literature to assess the global and regional patterns and impacts of illegal activities on species in protected areas and to provide information on the factors associated with population decline that may help improve the conservation of existing protected areas.

Anthropogenic threats reported from within PAs include hunting, logging, settlement, cultivation, livestock grazing etc. Each can reduce the ability of PAs to preserve biodiversity effectively (11). Moreover, the ability of PAs to protect species can be influenced by factors beyond their boundaries. For example, human-animal conflict has caused widespread mortality of carnivore species at and outside reserve borders, jeopardizing the persistence of populations inside protected areas, particularly for species with large home ranges that encompass both protected and unprotected land (12). At a larger scale, poaching of migratory species during periods when they are outside PAs has been reported as among the major threats imperiling the long-term conservation of species such as saiga (*Saiga tatarica*) and wildebeest (*Connochaetes taurinus*) antelopes, in Asia and East Africa respectively(13, 14). The ability of populations to sustain and recover from illegal activities within and surrounding PAs will depend on the type and level of activity, combined with the biological characteristics of the species affected by those activities. For example, selective poaching of males has been associated with the reproductive collapse and population decline of saiga (13), and species with low rates of reproduction and growth may be particularly prone to population decline and extinction (15).

While the studies above provide useful accounts of the threats facing PAs, relatively few of them quantify the relative contribution of individual threats to the overall pattern of population change and decline in PAs (16). Such an assessment is required to identify strategies to improve PA performance, such as where to target additional resources and which actions are most effective at enforcing existing regulations (17). Here, we review research published over 35 years (1980-1914) on illegal activities in PAs to understand the global and regional patterns and to assess their impacts on species in protected areas. We evaluate what factors determine the likelihood that illegal activities lead to the decline in the populations of targeted animals. In particular, we assess whether the different legal status in different PAs affects their ability to prevent population declines, and whether attributes of the species (i.e. body mass) and socio-economic context of a country (i.e. human development and agricultural land use change statuses) account for differences in PA success. Finally, we draw on these results to propose recommendations for reducing impacts of illegal activities in protected areas.

## Materials and Methods

### Data collation

We searched Web of Science, Science Direct, Google Scholar and Scopus for all papers since 1950 using the search terms: illegal activit*AND protected area OR region name (e.g. Europe, Asia, North/South America, Africa and Australia, New Zealand) OR reserve OR biodiversity outcome. Further search was performed incorporating poach / wildlife poaching AND protected areas OR reserve OR region name (as above). All online searches for the publications were conducted between 15^th^ November 2014 and 10^th^ March 2015. We screened the returns based on criteria (i-v below) that ensured the results from each paper were related to both PAs and illegal extraction of biodiversity;

i. Whether the research was done in a protected area and addressed issues of illegal extraction of biodiversity; animals, plants or both.
ii. Whether the research was on illegal activities by a human population and a mention was clear that the extraction is from the named protected area.
iii. If the research showed impact on species being extracted, decline or not declined
iv. Only used primary data papers not meta-analyses or reviews.
v. Studies covering similar sites and year of data collection were examined for relevance and only one that satisfied all criteria (i-iii) was included in the analysis.

We found no publications that fully satisfied the review criteria for the papers published between 1950 and 1980. Where two or more papers were published for the same protected area during a similar period of data collection, only one that satisfied all the criteria was used for the analysis. From each paper we extracted information on PA location, threat types, study species, perceived impacts of illegal resource use and purpose of research, continent (i.e. Africa, Asia, Europe, South America) and geographic region within continent (i.e. east, central, south and west). We recorded population trend (i.e. decline, no decline or unstated) for each PA/species combination from each paper as reported by the paper and the reasons mentioned for such outcome (e.g. illegal hunting, logging) to examine their effect on population status in the PA. For example, if a paper investigated protected areas X and Y and reported impacts of illegal activities on various species i, j, k…, then each species / site combination became one row in the dataset, including the impact scores (1= species decline or 0 = no decline or NA = no reported impact for that species) and any covariate information for PA or paper (author, year of study, PA name, etc.). The method arriving at the reported population trend status was recorded but not analyzed because most studies did not show the data used to arrive at a species outcome (e.g. decline), thus was difficult to tease apart whether the species decline was causal or correlative, a common problem in many meta-analysis studies (18).

Further, we extracted body mass data for each species. For mammals, we used the mammal database PanTHERIA (www.pantheria.org) and the current IUCN species red book data (19). We used body mass of closely-related species for two species that were not located in the mammal database (see supplementary material SM1). Body masses for reptile, amphibian and fish species were either extracted from the original papers (if provided) or from credible online material (supplementary material SM1). For bird body mass we used Dunning (20). To identify the geographical location of the study site, we cross-referenced the papers with the WCMC IUCN Protected Planet database (21) to identify the coordinates of the centroid of each PA. We also searched from this PA database for each name of the researched protected area and recorded its appropriate category under the current IUCN-PA categories.

To assess whether population change reported was related to wider scale economic or social change, we extracted country-level human development (HDI) and agricultural land use change (ALC) indices from the UNDP and World Bank databases (22, 23). ALC is an index of measure of the amount of land converted to agriculture and other human activities such as settlement. We calculated the ALC over a decade period encompassing the times when research for the reviewed papers were conducted as most papers did not report the exact dates of data collection. We used HDI as a predictor rather than measures of governance (the two are correlated) because HDI is a more direct indicator of development. Further, to understand the effect of different legal status on species decline we grouped the PAs into two levels of protection: strict PAs (for PAs under IUCN category I & II) and less strict PAs (categories III-VI). Furthermore, we categorized species into two broad groups of mammal and non-mammal for analysis to examine any differences between groups in the way they are threatened by illegal activities. We placed reptile, bird, fish and mollusc into one group: “non-mammal” as data were too sparse for each of these taxa to be tested individually in the model. There was no reported status for any of the plant species in the dataset; therefore, we excluded records of plants from the analysis.

### Data analysis

Several papers reported results for many species, and/or information for multiple protected areas. Here we refer to each unique combination of species within an individual PA (i.e. species × PA) reported in a paper as a study, with therefore potentially several studies per paper (see data extraction in methods). For modelling, we filtered all records where the population outcome was unknown. We analyzed spatial and temporal trends in research effort of the previous work on illegal activities and identified existing gaps. To assess whether population declines are associated with generic factors such as PA level of protection, type of poaching, species and a country’s socio-economic status, we used generalized linear mixed models (GLMM) with a binomial error term and logit link function using the statistical software R (version 3.2 R Development Core Team, 2015).

We built an initial global model incorporating seven fixed factors: human development index (HDI), percentage land use change index (ALC), log species body mass, poaching type (commercial, subsistence, or a mixture), PA level of protection (i.e. IUCN categories classified as strict and less strict), continent (Africa/Asia/America/Europe) and species group (i.e. mammal and non-mammal). Because different species could relate to the same PA and country as studies from other papers at different times, we accounted for this by including country, paper and PA as random effects in all models and fitted the datasets using the ‘glmer’ function implemented in the R-package ‘lme4’(24). We used a backwards stepwise removal of non-significant terms (with Chi-test) to evaluate the relative effect of each factor on the population decline. We obtained model confidence intervals around variables showing statistical significance in the minimum adequate model using the Wald-method (24). Furthermore, in each model we evaluated whether the likelihood of finding an impact on a species’ population was due to the PA level of protection and level of hunting (subsistence, commercial or both) and species group and geographic regions. We also examined whether log species body size, country’s human development index and agricultural land use change index were consistently correlated with the population decline.

African elephants (i.e. mainly the African savanna elephant *Loxodonta africana*, but with some data for African forest elephant, *Loxodonta cyclotis*; no data on Indian elephants were included in the dataset), hereafter referred to as ‘elephant’, had large numbers of studies in the dataset (contributing 22 % of all records), probably reflecting the increasing concerns for its poaching (25). To check the generality of our results, we repeated the analysis using two subsets of the data: once for elephants alone, and once for all animal species except African elephants. Similarly, because there were sufficient data to analyze some components separately we repeated all the analyses with and without the strict PAs (IUCN categories I&II) as well as on separate subsets of data including Africa and Asia continents to examine their impact on the whole data set and the relative effects of different predictors between the two continents.

## Results

### Spatial distribution and trends in research on poaching in PAs

We identified 1598 papers that met the initial search criteria, of which 92 (Supplementary material SM2: reporting 1048 species x PA results = ‘studies’) met all the requirements and were used for analysis. The 92 published papers researched 146 protected areas from four continents, with the largest number of studies from Africa and Madagascar (819 studies), followed by Asia (162), South and Central America (66), and Europe (1) (Figure 1).

**Figure 1.**
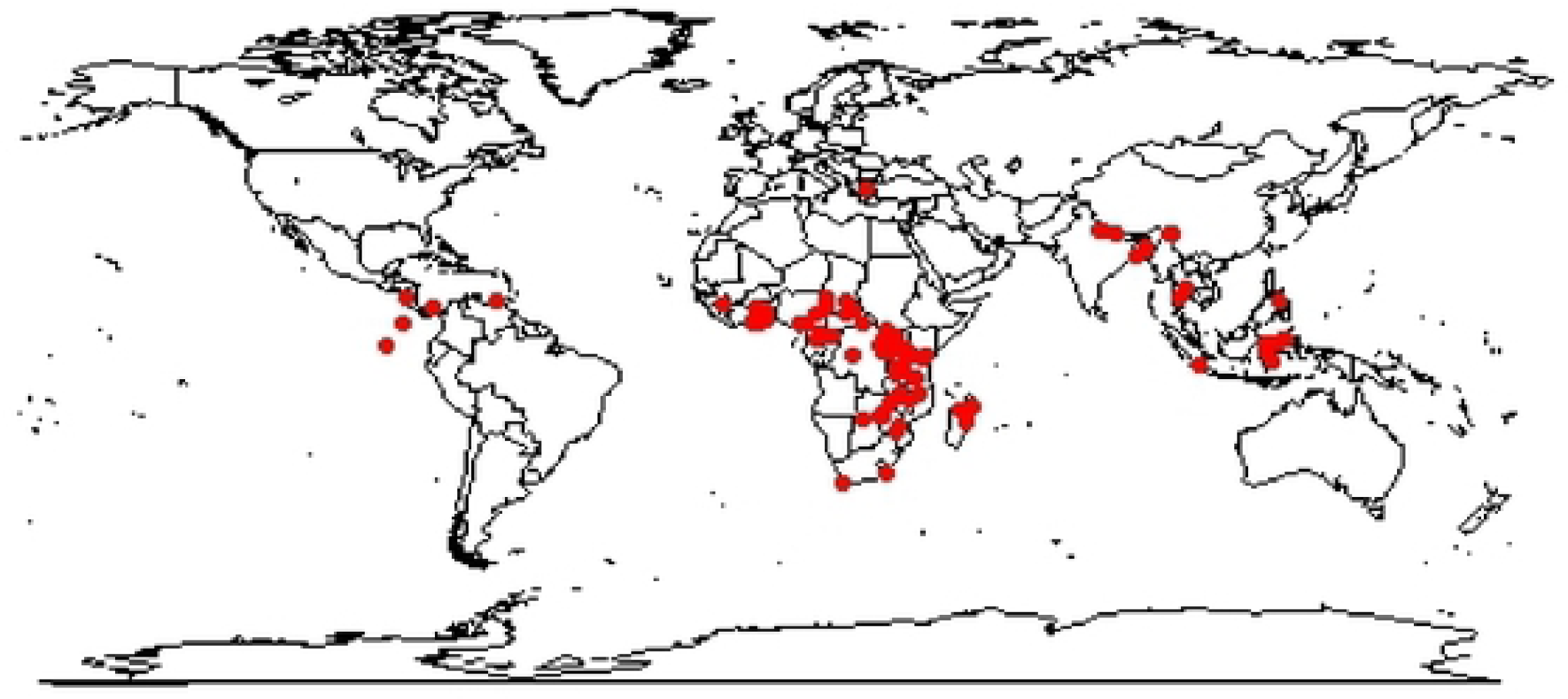
The locations of all PAs investigated (red dots correspond to centroids of protected areas). Numbers of studies (species x PA combinations) were: Africa and Madagascar (819), Asia (162), South and Central America (66), and Europe (1).

Most papers focused on single PA (i.e. local scale, n = 54 papers), or few PAs existing as one contiguous ecosystem and landscape (n = 39 papers). All protected area types were investigated but the IUCN category II level of protection was researched the most (57.35%, n = 65 papers; Figure 2). There was no paper published between 1950 and 1979 that satisfied our search criteria: all relevant papers were published in 1980 onwards.

**Figure 2.**
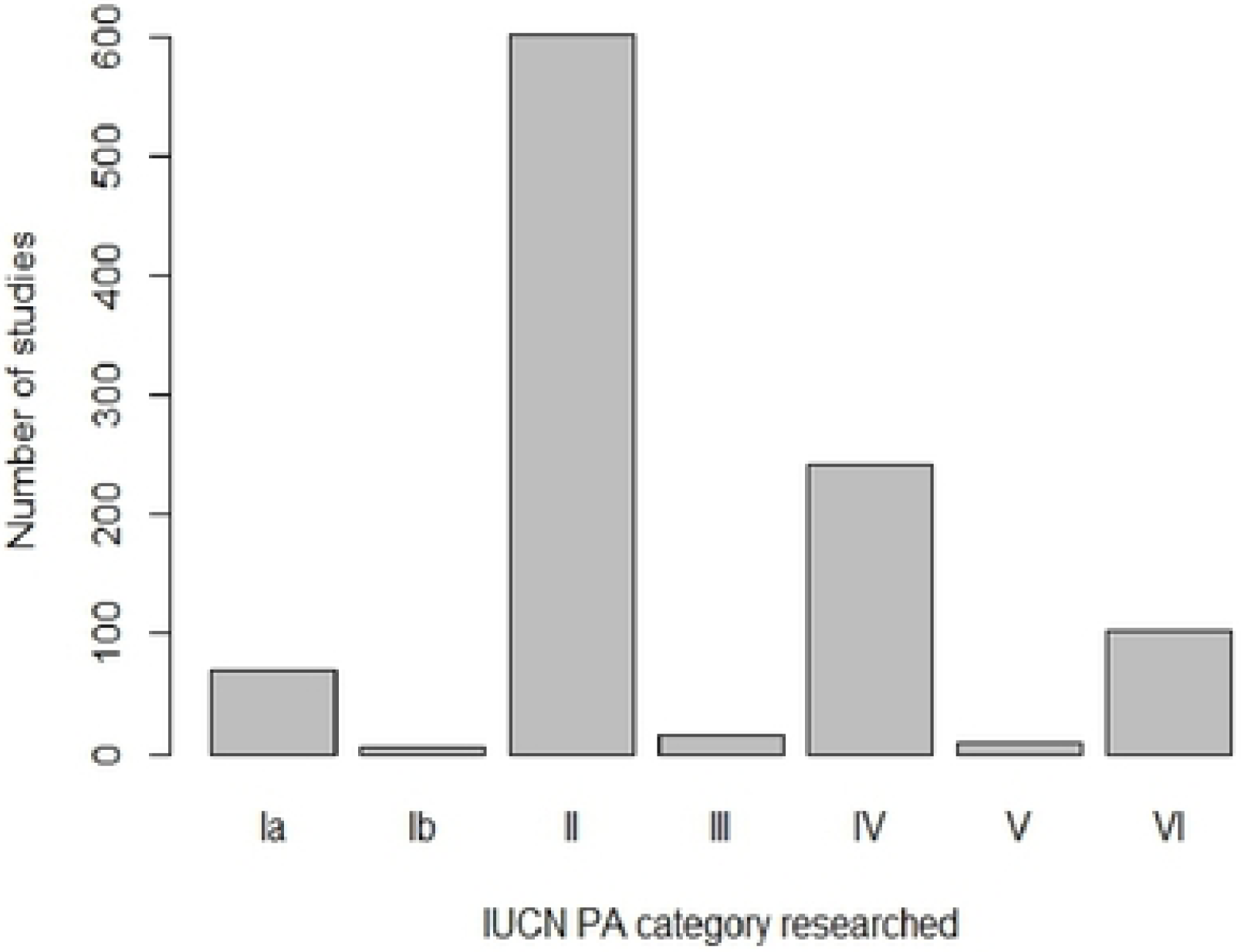
The number of studies (species x PA combinations) on illegal activities carried out in PAs with different levels of IUCN protection (I, greatest protection; VI, least). This is based on the 92 papers published between 1980 and 2014, encompassing four continents. There is a strong bias towards the PA category II. The IUCN PA categories are used to facilitate comparisons of different PA ‘entities’ in different countries (e.g. conservation area, forest reserve, game-controlled area, game reserve, game sanctuary, marine reserve, national park and nature reserve).

The research had varying purposes: investigating impacts (71 papers); conservation rationale (e.g. providing new methods for investigating illegal activities; 17 papers) and management/control of illegal activities (5 papers). Further, research has increased substantially during the last 35 years with greater number of published papers since 2005 (Figure S1); most of this increase was in Africa.

Furthermore, we found a temporal increase in the variety of research methods throughout the period (Figure 3). Interviews (n = 22 papers), animal counts (n = 28) and patrols (n = 9) were the most commonly used methods in the literature. Several research projects (n = 34) combined methods, while two used snare surveys. Animal count was the dominant research method during the first decade (1980-1990) and has increased in use since then. Use of interviewing and patrolling methods to investigate illegal activities in PAs has been used between 1990 and the present (i.e. 2014).

**Figure 3.**
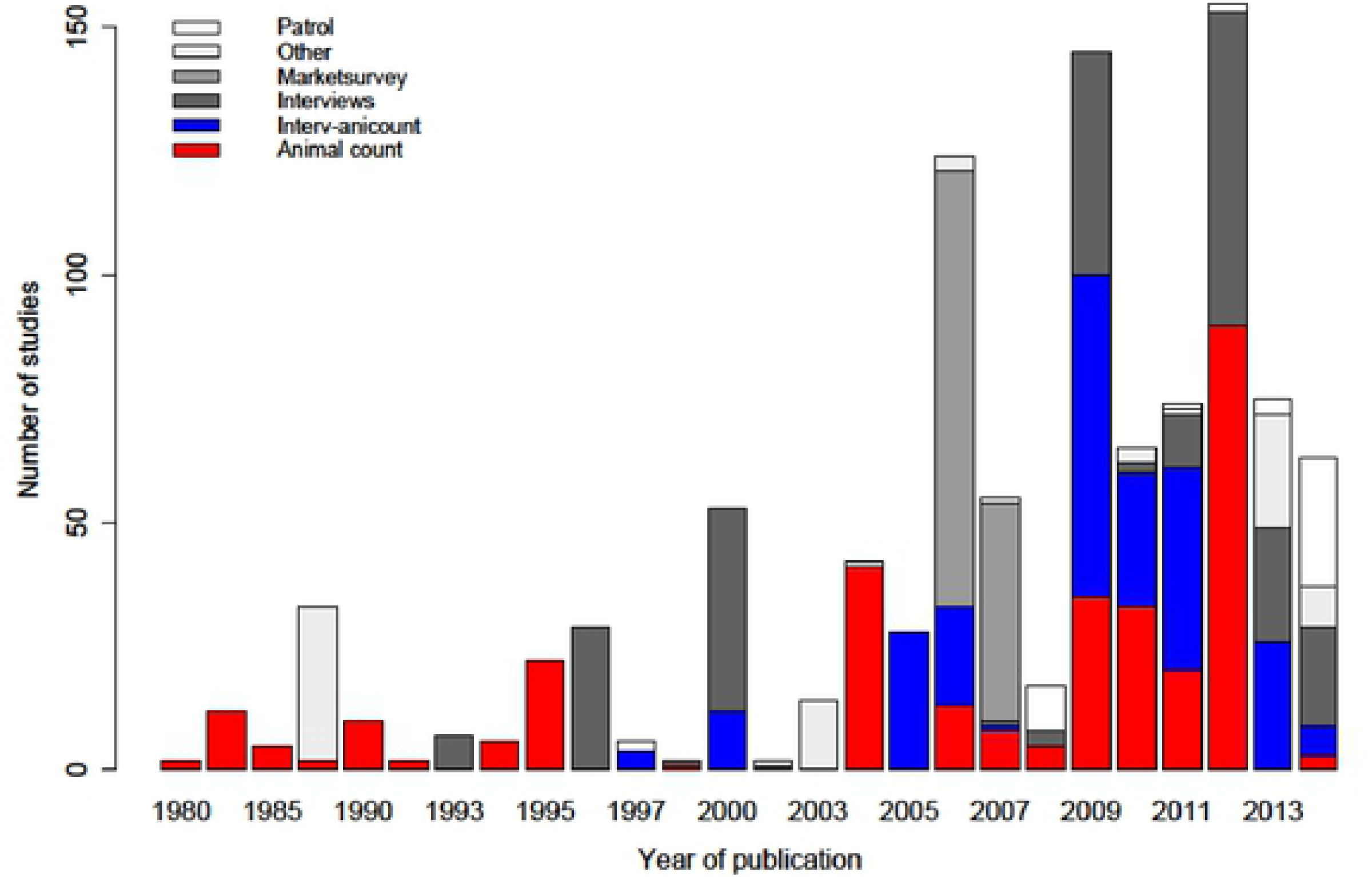
Methods used in the 92 publications considered. Recent studies show an increasing diversity of methodological approaches. ‘Other’ includes bone collection from poacher camps and sporadic field observations.

### Patterns of species extraction and country socio-economic status

Three hundred and fifty-three species, comprising mammals (220), reptiles (18), birds (17), fishes (12), molluscs (2) and plants (84) were reportedly harvested illegally in the 146 protected areas (Table S1). These species were extracted for subsistence use (10.8%, N = 221), commercial use (19.9%) or both (60.6%). Almost nine percent of the studies did not report a reason of illegal resource extraction, and none of the plant studies included all the data that were required for this analysis. The countries where research was conducted showed varying levels of development (Figure 4) and land use change (Figure 5).

**Figure 4.**
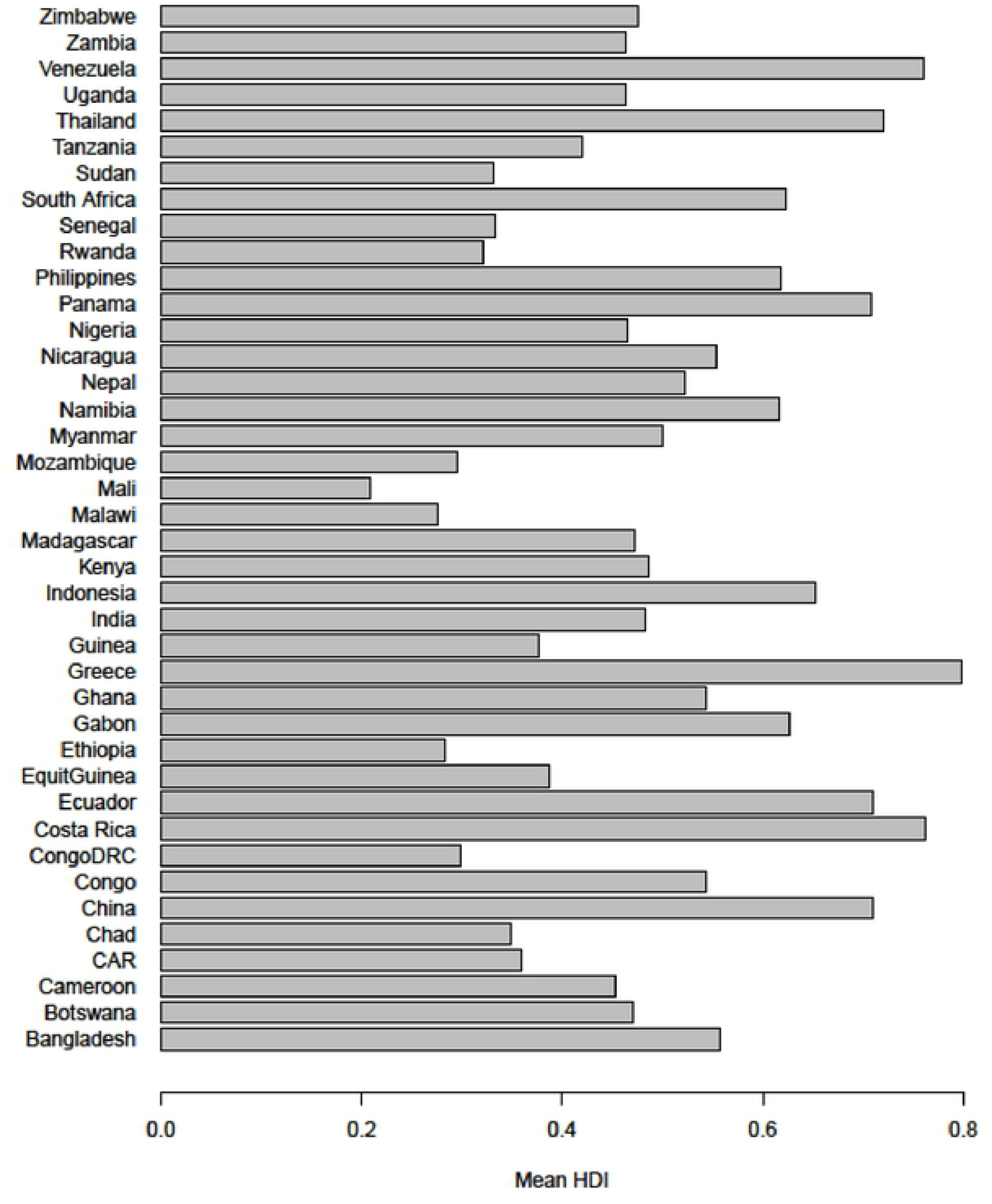
Human development index (HDI) for the countries where research was conducted. HDI values shown are the mean scores for the total number of years that a country was researched.

**Figure 5.**
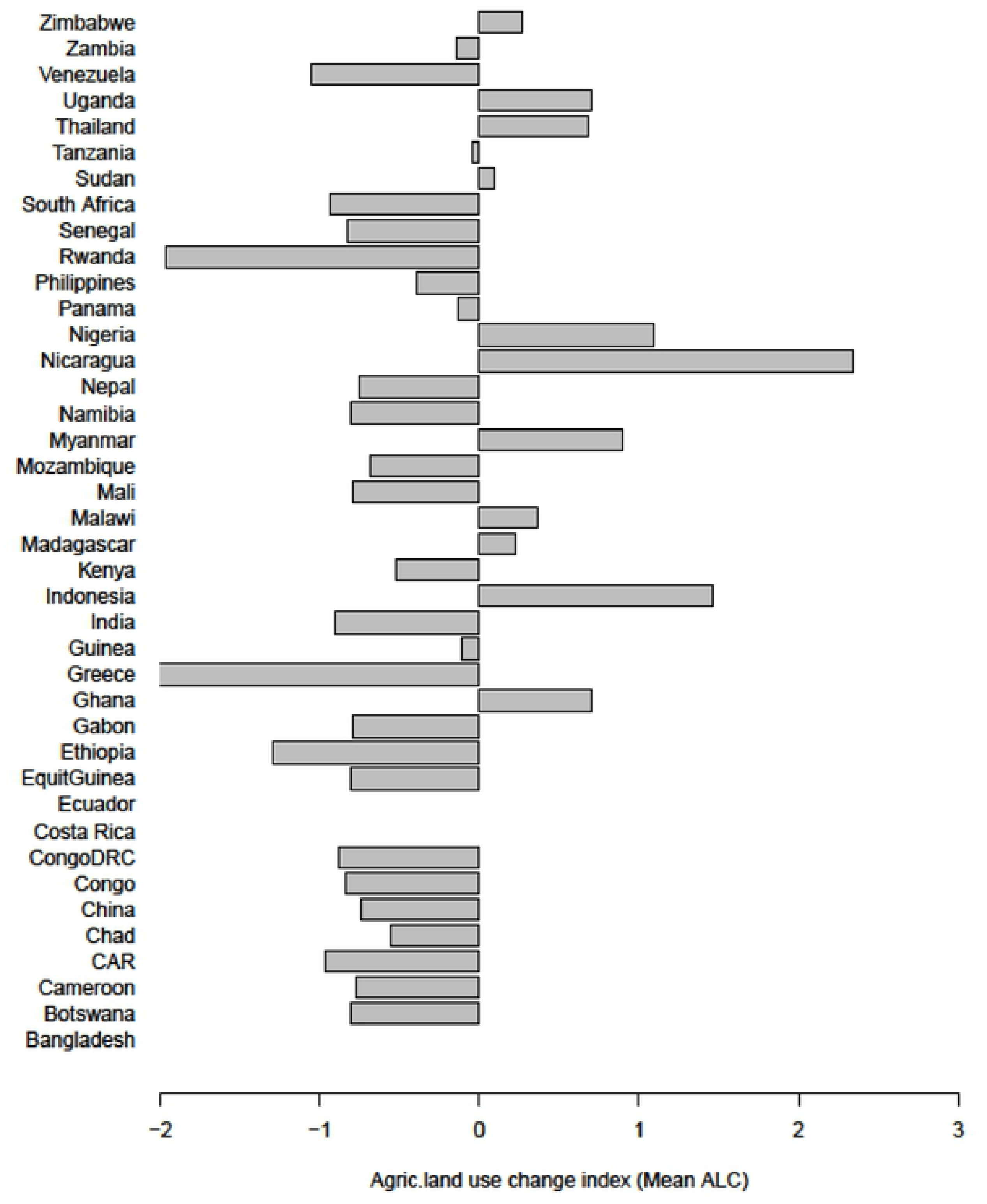
Agricultural land use (ALC) change index of a country where research was conducted. Negative change infers to loss of the natural habitats to agricultural activities and settlement. Mean ALC includes the total number of years a particular country PAs were researched during the last 35 years.

### Socio-ecological and geographic factors influencing population decline in PAs threatened with illegal activities

Species body mass and species group (mammal or non-mammal) strongly influenced probability of species decline in all PA types with species with greater body mass especially mammals experiencing larger population decline (Model 1 in **Error! Reference source not found.**, Figure 6a). There was an effect of the dominant species in our dataset. When removed the African elephants, we found that species declines were greater in PAs faced with commercial rather than subsistence poaching alone (Figure 6b); as before, mammals and species with greater body mass also exhibited the greatest declines in these models (Model 2 in **Error! Reference source not found.**).

**Figure 6.**
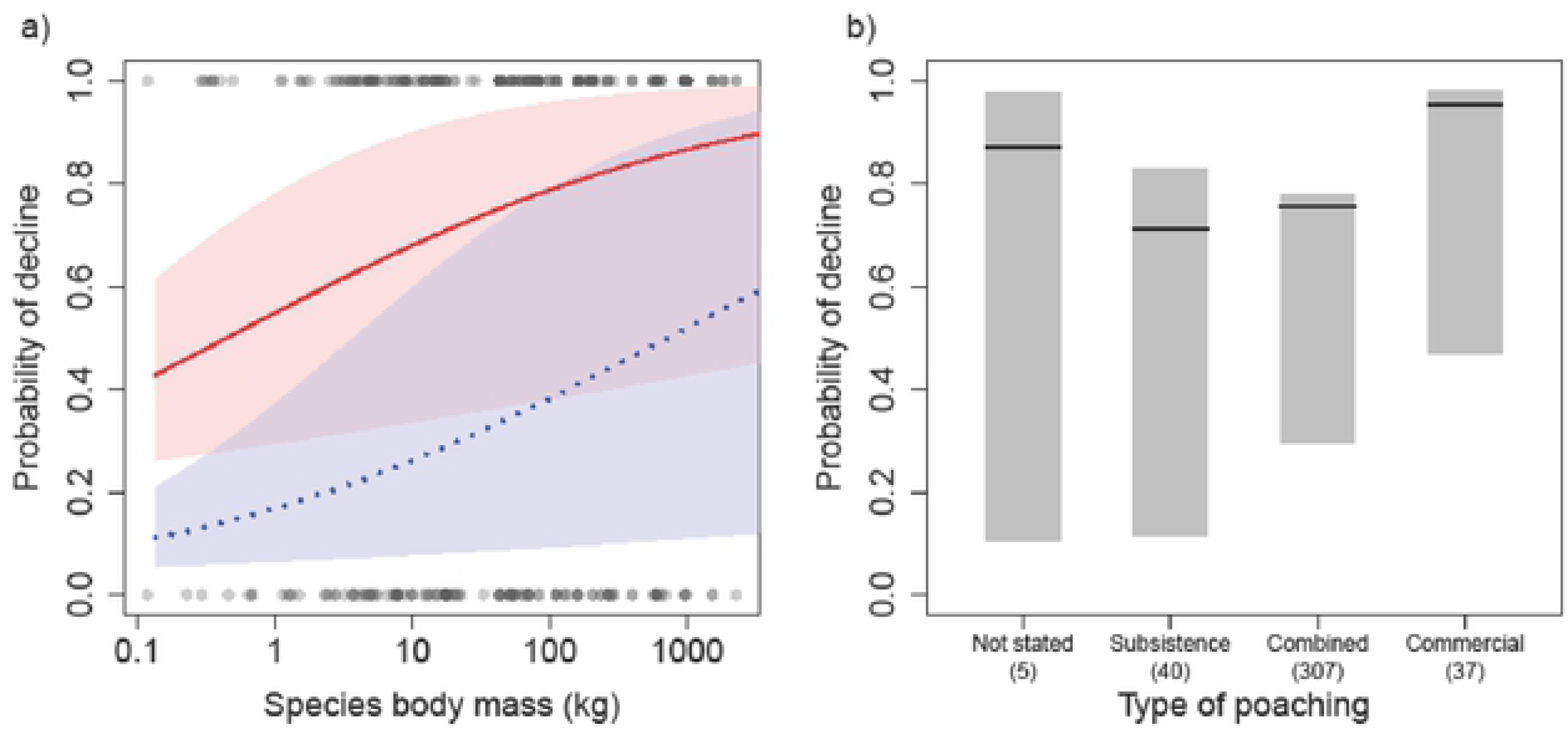
Effect of species body mass (a) and level of poaching (b) on the probability of decline of animal species (mammal-upper solid line, and non-mammal-lower dashed line) excluding the dominant species (African elephants) across all PAs (IUCN I-VI). Large bodied species had higher risk of decline when threatened with illegal activities. Shaded area shows 95% CI around the estimates of effect size for the mammal and non-mammal species.

Separate analysis on PAs according to protection level revealed variable results. Human development index (HDI), species body mass and species groups strongly influenced the probability that species would decline in strict PAs (Model 3 in Table 1, Figure 7). Strict PAs in low human development index countries were associated with increased species decline, of mammal species with greater body mass. In contrast, in less strict PAs (IUCN category III-VI), we found species decline was best explained by agricultural land use change (ALC) in the wider area; i.e. species in all PAs with greater habitat loss were more likely to have experienced population decline (Model 4 in Table 1).

**Table 1.**
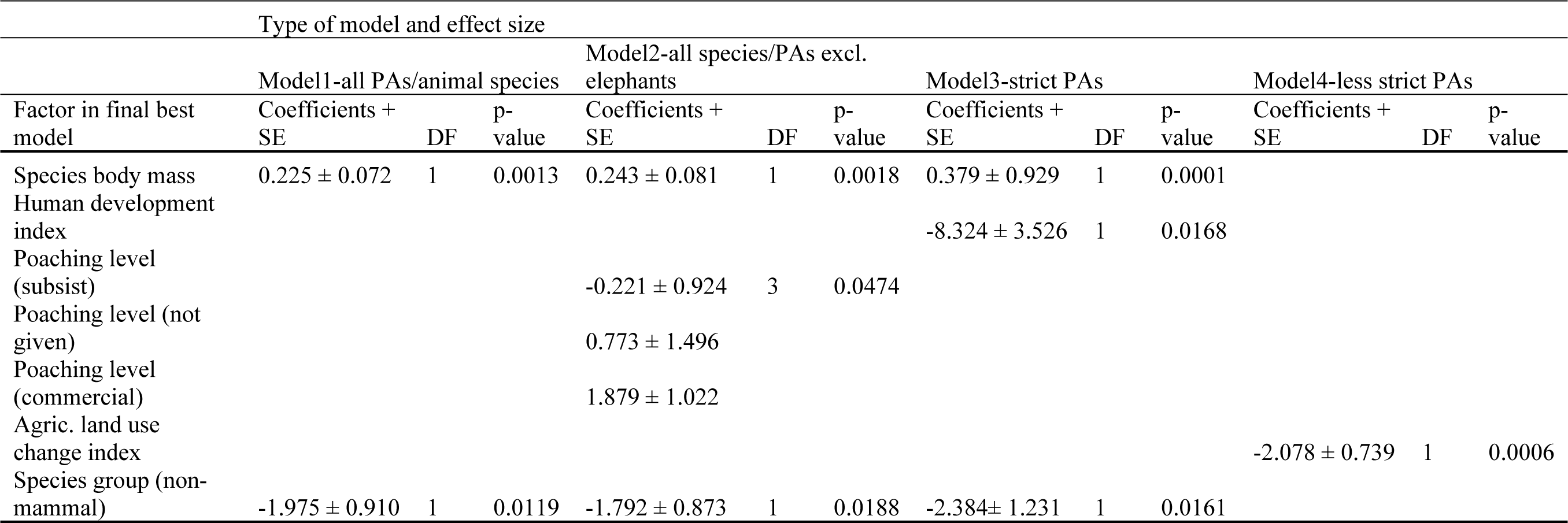
Results of the GLMM best models showing probability of species decline and the explanatory factors in PAs across four continents. Coefficient sign (+/−) indicates size of effect of covariables i.e. increase for plus and decrease for minus.

| Type of model and effect size |  |  |  |  |  |  |  |  |  |  |  |  |
| --- | --- | --- | --- | --- | --- | --- | --- | --- | --- | --- | --- | --- |
| Factor in final best model | Model1-all PAs/animal species |  |  | Model2-all species/PAs excl. elephants |  |  | Model3-strict PAs |  |  | Model4-less strict PAs |  |  |
|  | Coefficients + SE | DF | p-value | Coefficients + SE | DF | p-value | Coefficients + SE | DF | p-value | Coefficients + SE | DF | p-value |
| Species body mass | 0.225 ± 0.072 | 1 | 0.0013 | 0.243 ± 0.081 | 1 | 0.0018 | 0.379 ± 0.929 | 1 | 0.0001 |  |  |  |
| Human development index |  |  |  |  |  |  | -8.324 ± 3.526 | 1 | 0.0168 |  |  |  |
| Poaching level (subsist) |  |  |  | -0.221 ± 0.924 | 3 | 0.0474 |  |  |  |  |  |  |
| Poaching level (not given) |  |  |  | 0.773 ± 1.496 |  |  |  |  |  |  |  |  |
| Poaching level (commercial) |  |  |  | 1.879 ± 1.022 |  |  |  |  |  |  |  |  |
| Agric. land use change index |  |  |  |  |  |  |  |  |  | -2.078 ± 0.739 | 1 | 0.0006 |
| Species group (non-mammal) | -1.975 ± 0.910 | 1 | 0.0119 | -1.792 ± 0.873 | 1 | 0.0188 | -2.384 ± 1.231 | 1 | 0.0161 |  |  |  |

**Figure 7.**
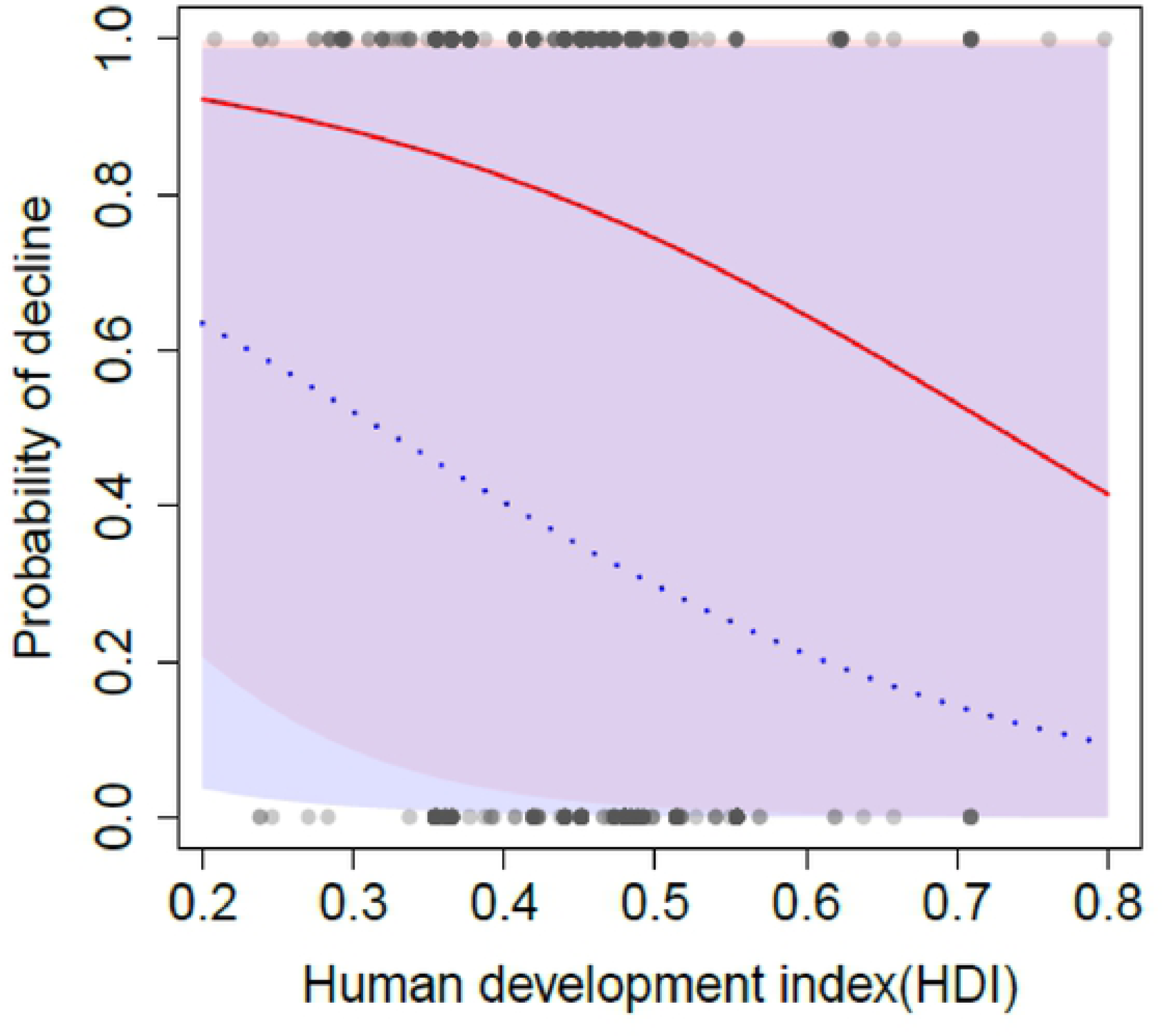
HDI influence on population decline of animal species (upper solid line and lower dotted line for mammal and non-mammal respectively) within strict PAs (IUCN category I&II). Least developed countries had the highest probability of their PAs experiencing species decline. Shaded areas show 95% CI around the estimates of effect size of this factor.

The impact of illegal activities on African elephants was variable. Across all PAs (IUCN I-VI), decline was highly associated with lower human development index countries, high habitat loss; i.e. negative agricultural land use change index and with the geographic region of PA location (Model-All PAs in **Error! Reference source not found.**, Figure 8). This decline was greatest in central Africa, followed by the east and west; and least in southern Africa. In an analysis of strict PAs alone we found two factors (i.e. human development index and agricultural land use change index) associated with the increase in elephant decline (Model-strict PAs in Table 2). On the other hand, we found geographic region significantly associated with increased elephant decline within less strict PAs in central and east Africa (Model-less strict PAs in **Error! Reference source not found.**).

**Table 2.** Results from GLMM models with various factors explaining the probability of decline of African elephants across all PA types and separately between strict and less strict PAs.

| Factor in final best model | Type of model and effect size |  |  |  |  |  |  |  |  |
| --- | --- | --- | --- | --- | --- | --- | --- | --- | --- |
|  | All PAs types (cat. I-VI) |  |  | Strict PAs (cat. I & II) |  |  | Less strict PAs (cat. III-VI) |  |  |
|  | Coefficients +<br>SE | DF | p-value | Coefficients +<br>SE | DF | p-value | Coefficients +<br>SE | DF | p-value |
| Human development index | -12.350 ± 4.207 | 1 | 0.0005 | -19.459 ± 6.496 | 1 | 0.0004 |  |  |  |
| Agri. land use change index | -1.461 ± 0.699 | 1 | 0.0116 | -1.173 ± 0.826 | 1 | 0.0371 |  |  |  |
| Africa zone (East) | 0.723 ± 1.116 | 3 | 0.0100 |  |  |  | 0.406 ± 1.307 | 3 | 0.0282 |
| Africa zone (South) | -2.186 ± 1.041 |  |  |  |  |  | -2.773 ± 1.275 |  |  |
| Africa zone (West) | 0.075 ± 2.749 |  |  |  |  |  | 0.981 ± 0.677 |  |  |

**Figure 8.**
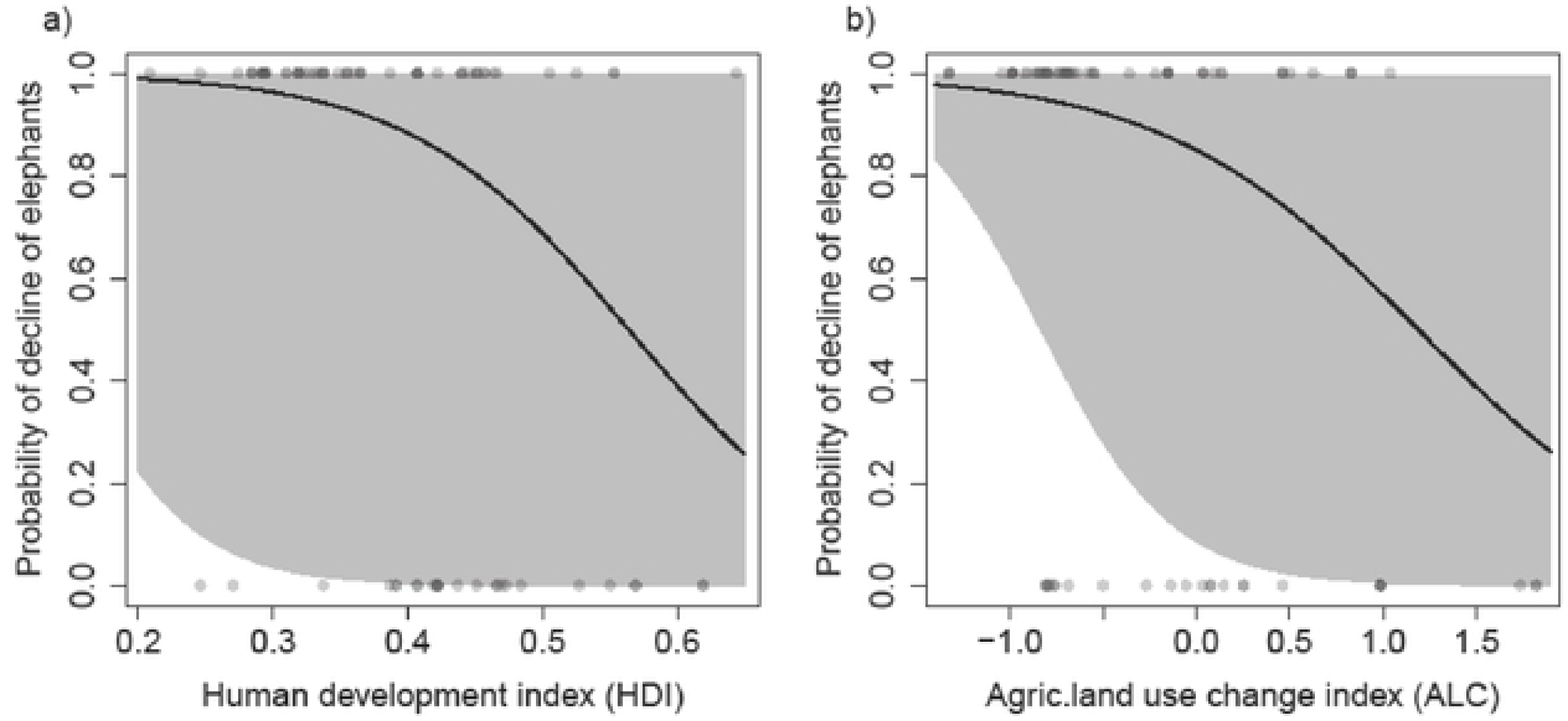
The country’s human development index (a) and agricultural land use change index (b) as best predictors of decline in the African elephant within in all PAs (category I-VI) across the African continent. High negative ALC values are associated with severe habitat loss and a high risk of population decline of African elephants. Gray area shows 95% CI around the estimates in the minimum adequate GLMM model.

Furthermore, the probability that illegal activities were associated with population declines also varied regionally across PAs in Africa, Asia and Latin America. Lower human development index of countries, greater species body mass and species group (i.e. mammal) were the strongest predictors of increased species decline in PAs across Africa (Model -Africa in **Error! Reference source not found.**). In contrast, species decline in Asia was strongly positively associated with PA strictness (Model-Asia in Table 3). On the combined data for Asia and Latin America, we found that increased probability of species decline in PAs across these continents was correlated with greater agricultural land use change (Model combined in **Error! Reference source not found.**).

**Table 3.** Results from GLMM models with influence of the various factors on the probability of species decline in PAs separately for Africa continents and on combined data for Asia and Latin America.

| Type of model and effect size |  |  |  |  |  |  |  |  |  |
| --- | --- | --- | --- | --- | --- | --- | --- | --- | --- |
| Factor in final best model | Africa-all animals/PAs |  |  | Asia-all animals/PAs |  |  | Asia + L. America |  |  |
|  | Coefficients + SE | DF | p-value | Coefficients + SE | DF | p-value | Coefficients + SE | DF | p-value |
| Human development index | -8.657 ± 3.708 | 1 | 0.00386 |  |  |  |  |  |  |
| Body mass | 0.338 ± 0.084 | 1 | 0.00022 |  |  |  |  |  |  |
| Species group (non-mammal) | -2.269 ± 1.095 | 1 | 0.01223 |  |  |  |  |  |  |
| PA strictness |  |  |  | 1.974 ± 0.684 | 1 | 0.0237 |  |  |  |
| Agric. land use change index |  |  |  |  |  |  | -13.946 ± 4.741 | 1 | 0.0077 |

## Discussion

Biodiversity in protected areas faces numerous anthropogenic threats (11). We analyzed data published since the 1980’s to understand impacts of illegal activities on species decline in protected areas. There were a strong taxonomic and geographic biases in research on illegal activities within protected areas during the review period. We found that population declines were more likely consequences of illegal activities in countries with low human development index (HDI). Different groups of species declined at different rates, with large bodied mammals mostly likely to show population decline. As well as poaching, we also found that habitat loss had an additional impact on population decline of animal species in less strict PAs, particularly in countries experiencing greater agricultural land use change. Further, species decline was also associated with geographic region of PA location being greater in Africa than Asia or Latin America.

The identification of correlations between human development indices and illegal activities in this study supports a widely held view, e.g. Bennett, Blencowe (26), Adams, Aveling (27), but one that is often based on limited data (28): that biodiversity decline is greater in relatively poor regions. Low human development scores could impact illegal activities in two ways: firstly, poor people may tend to exploit species illegally from PAs because they have limited alternatives (29). Secondly, poor countries have fewer resources to invest in PA conservation. Underfunding may result in increased illegal activities in PAs due to insufficient law enforcement (30). This is supported by our model encompassing the most strictly protected areas alone, which indicates high probability of species decline associated with low human development index countries (Table 1). Hilborn, Arcese (31) demonstrated that increased funding budgets for anti-poaching activities in the Serengeti National park greatly reduced poaching pressures and lead to the recovery of the buffalo population.

However, increasing conservation funding may not necessarily result into improved conservation particularly when social and political constraints exist. For example, social and political unrest may increase rates of illegal activities, reduce wildlife populations and thwart conservation efforts altogether (32, 33). Our results provide evidence that poverty, in as much as it is measured by the HDI, may have significant negative impacts on species due to accelerated poaching, whether that be because of increased external pressures on PAs or decreased policing and protection within strict PAs. Further, illegal activities increased species vulnerability to decline through increased habitat loss mostly in less strictly protected areas (Table 1). This could be because these PAs are often afforded minimal protection (34) and therefore this exposes them to intense illegal activities making them less reliable for the effective conservation of large and medium-size mammals (35). These findings also provide evidence that supports Craigie et al.’s (2010) assertions that illegal hunting and habitat loss are the major causes of continental-wide declines in megafauna in PAs across Africa. Our study highlights the need to consider human development issues more seriously to ensure effective conservation of biodiversity within existing protected areas.

Large bodied species are likely highly susceptible to decline because they have slow growth rates and so overharvesting is likely to cause population decline (36). Low population growth rates in combination with multiple threats from poaching and diseases are known to have significant impact on population persistence (12, 15). By contrast, smaller mammals (with higher reproductive and growth rates) showed fewer declines and appeared to sustain harvest, though relatively few small species are the specific targets of poaching in the PAs. Our model that excluded elephant suggested that commercial poaching has the greatest potential to cause population decline. Further, the propensity for elephants to decline in low human development index countries and geographic regions, and where habitat loss was taking place (Table 3), is consistent with recent analyses that this species is threatened with poaching and habitat loss across its habitat range in Africa (37, 38).

The pattern of species declines across the network of protected areas is worrying and suggests that PA policing (including access to appropriate conservation information) and resources need to be improved. PA-specific information is important for understanding how illegal activities vary spatially and across time and there is a need to be able to predict future trends and thereby possible future management strategies e.g. Critchlow, Plumptre (39). Furthermore, land use change poses additional risks of species decline in PAs across Asia and Latin America, relative to Africa, and its impact was more severe in less strict than strict PAs (**Error! Reference source not found.**). This could be attributable to the habitat loss and poaching occurring inside these protected (40–42), or the wider effects on animals that roam outside protected areas for parts of the year. These results are consistent with previous studies that have reported biodiversity decline and loss within PAs in these regions e.g. Geldmann, Barnes (16), Harrison (43), Laurance, Carolina Useche (44). Our findings suggest that land use change is a major threat that requires urgent attention to improve PA conservation across Latin America and continental Asia.

The geographical bias in the spatial distribution of research observed in these data is likely a consequence of interests among the researchers rather than being driven solely by the levels of illegal activities in particular PAs or countries. However, the temporal and spatial patterns of research observed in this study provide insight into the extent of the problem of illegal activities in PAs and therefore suggest that PAs are currently in need of new strategies to minimize impacts of illegal activities and to improve their conservation effectiveness (45). To date, research effort has concentrated on quantifying the extent and impact of illegal activities on focal species; in other words, documenting the problem. Far less information is available on which conservation management strategies (including human development and preventing illegal international trade, as well as within-PA activities) are most effective at reducing illegal activities. Only 5 out of 92 publications reviewed here explicitly considered how the management of illegal activities might affect population declines. New research should focus on developing and testing new methods for reducing levels and impacts of illegal activities on species in PAs.

## Conclusion

Tackling illegal activities within protected areas remains a high conservation priority. Our results suggest that a combination of strategies may be required that simultaneously reduce the frequency with which illegal activities are attempted and reduce the likelihood that such attempts will be successful (from the perspective of the perpetrators). Regarding the former, we found that illegal activities in poor countries often lead to population declines within PAs. These findings suggest that poverty alleviation may be an appropriate conservation strategy to reduce illegal activity pressures (26). The implication of this for local and national policies is that more effort needs to be invested to improve the social and economic status of the human populations. This needs to work in tandem with increasing the effectiveness of traditional conservation activities to prevent illegal activities; which may itself reduce the inclination of people to attempt future illegal activities. At the international level, these results may imply that PAs in low HDI countries may need more international support to curb pressures of illegal activities, particularly those driven by trans-boundary forces. Sound strategies to stop elephant poaching and ivory trade, and trade in bushmeat, as well as strengthening collaboration in conservation and research, are necessary to improve PA effectiveness in these countries.

## Acknowledgements

We are grateful to the anonymous reviewers for the comments on earlier drafts of this manuscript. AAR is a commonwealth scholar funded by the U.K. government.

## SUPPORTING INFORMATION CAPTIONS

Table S1. The reasons recorded by the studies to explain the status of biodiversity reported in the reviewed papers. Plus (+) sign indicates the reason was mentioned together with the poaching in the PA. Studies (43.2%, n = 453) that sum up to the totals did not indicate the status of species being investigated and are not shown here.

Figure S1. Increasing publication trend for illegal activities in PAs during the last 35 years.

Table S2. 353 species extracted and threatened with illegal activities in 146 PAs across four continents as published in the last 35 years (1980-2014)

SM1: List of Additional Sources of data for Species body mass

SM2: Bibliography of the 92 research papers used in this analysis

